# Deep-learning based bioactive therapeutic peptides generation and screening

**DOI:** 10.1101/2022.11.14.516530

**Authors:** Haiping Zhang, Konda Mani Saravanan, Yanjie Wei, Yang Jiao, Yang Yang, Yi Pan, Xuli Wu, John Z.H. Zhang

## Abstract

Many bioactive peptides demonstrated therapeutic effects over-complicated diseases, such as antiviral, antibacterial, anticancer, *etc*. Similar to the generating *de novo* chemical compounds, with the accumulated bioactive peptides as a training set, it is possible to generate abundant potential bioactive peptides with deep learning. Such techniques would be significant for drug development since peptides are much easier and cheaper to synthesize than compounds. However, there are very few deep learning-based peptide generating models. Here, we have created an LSTM model (named LSTM_Pep) to generate *de novo* peptides and finetune learning to generate *de novo* peptides with certain potential therapeutic effects. Remarkably, the Antimicrobial Peptide Database has fully utilized in this work to generate various kinds of potential active *de novo* peptide. We proposed a pipeline for screening those generated peptides for a given target, and use Main protease of SARS-COV-2 as concept-of-proof example. Moreover, we have developed a deep learning-based protein-peptide prediction model (named DeepPep) for fast screening the generated peptides for the given targets. Together with the generating model, we have demonstrated iteratively finetune training, generating and screening peptides for higher predicted binding affinity peptides can be achieved. Our work sheds light on to the development of deep learning-based methods and pipelines to effectively generating and getting bioactive peptides with a specific therapeutic effect, and showcases how artificial intelligence can help discover *de novo* bioactive peptides that can bind to a particular target.

## Introduction

Many bioactive peptides have therapeutic effects against various complex diseases, such as antiviral, antibacterial, and anticancer[1]. More than 80 peptide drugs are currently approved to treat various diseases, including diabetes, cancer, osteoporosis, multiple sclerosis, HIV infection, and chronic pain[2–4]. The development of peptide synthesis methods[5] and the advancement of peptide drug delivery system technology[6,7] have also extensively promoted the development of peptide drugs. However, there is an extremely high diversity of amino acid sequences; for example, a peptide composed of standard amino acids with a length of 8 has theoretically 20^8 kinds of possible peptide sequences. If searching for potential peptides with lengths longer than eight by ergodic process, it is far beyond the current computing resources. Most research methods are based on local mutation of existing active peptides[8–10], while the diversity of such mutated peptides is often insufficient. Furthermore, it is challenging to find peptides with stronger activity or different binding mechanisms.

Peptide generation mainly relies on random mutation or optimizing existing peptides. Recently, there are some machine learning or deep learning-based models for peptide generation. A DeepImmuno-GAN architecture was developed to generate potential peptides binding to MHC[11]. Codon-Based Genetic Algorithm (CB-GA) method was proposed to generate de novo compounds[12]. Other deep learning-based methods for generating potential active peptides also have developed, including PepVAE[13], ProteinGAN[14], HydrAMP[15], PepGAN[16], peptide VAE[17]. Interestingly, Nagarajan et al. have used LSTM model to predict MIC value of peptide (the lowest concentration of an antibiotic at which bacterial growth is completely inhibited peptide). To our knowledge, there still no LSTM based model for peptide generation and finetuning to obtain *de novo* active peptide. However, there are much more existing small-molecule generative models for small molecules [18–20], and small molecules with specific potential biological activities can be targeted by finetuning. Similar to the generation of *de novo* compounds, with the increasing accumulation of active biological peptides as a training set, it is now possible to use deep learning models to generate many potentially biologically active peptides. Interestingly, it is expected that through iterative generative screening, we will eventually obtain biological activities that are much higher than known activities, such as super antimicrobial peptides[21]. Similar technologies will significantly facilitate drug development, as peptides are easier to synthesize than compounds and cheaper to purchase. Developing a deep learning model to generate peptides may help ensure the diversity of generated peptides. At the same time, through finetuning learning of peptide datasets with specific functions, a large number of peptides with potential specified activities can be generated, and sufficient diversity is ensured to find different sequences and structures of active peptides. Many designated active peptides are generated, which is far less than 20^8 and enables the fast identify active peptides that act on specific targets through subsequent screening methods.

Peptide protein docking can use public software, such as rosetta[22], ZDOCK[23], *etc*., but few comprehensive and efficient screening pipelines exist. Rosetta software is widely used among the existing protein-peptide binding prediction and design methods. Still, this method is slow, and it isn’t easy to screen out peptides with ultra-high affinity to the specified target protein in a short period. However, it is worth noting that the development of rosetta, modeling, molecular dynamics simulation, and other methods has extensively promoted the advanced screening of new peptide drugs. Recently, some deep learning based protein-peptide model also developed to prediction potential bioactive peptides or predict protein-peptide interaction[24–26]. Among them, the DeepACP use deep learning algorithm to identify anticancer peptides(ACPs)[24]. They tried convolutional, recurrent, and convolutional-recurrent networks to distinguish ACPs from non-ACPs, and found recurrent neural network with bidirectional long short-term memory cells is superior to other architectures. A deep learning method called XDeep-AcPEP was also developed for anticancer peptide activity prediction based on convolutional neural network and multitask learning[26]. Yipin Lei, et al. have proposed a deep-learning framework called CAMP for binary peptide-protein interaction prediction and peptide binding residue identification[25].

Small compounds screening have developed fast due to the fast development of deep learning[27,28] and MD simulation method[29] in protein-ligand interaction prediction. According to our previously work[30,31], utilize multiple different methods to screening drug compounds would have an advantage. Since protein-compound interaction have share many similarities with protein-peptide interaction, such hybrid screening strategy can provide insight for efficient and accurate identifying peptides for a given target.

In the present work, we created an Long Short-Term Memory (LSTM) model[32] for generating *de novo* peptides and then finetuning the model by training over known bioactive peptides to generate novel peptides potentially with same therapeutic activity. Furthermore, we developed a protein-peptide prediction method to screen the generated large number of potentially active peptides for a given protein target. We built screening pipeline by integrating deep learning, docking and MD simulation methods. The specific process is 1. generate potential peptides with specified activity, 2. Use tr-rosetta[33] to build a 3D structural model of peptides based on peptide sequences, 3. Then use ZDOCK to dock the peptides to specific pockets of given targets,4. Then use rosetta to perform flexible docking (or use Protein-peptide deep learning prediction to obtain high confidential conformation from ZDOCK result),5. Molecular dynamics simulations,6. Metadynamics simulations are used to evaluate binding free energy surfaces. Finally, a high-efficiency iteratively generate and screening strategy, which depend on LSTM_pep and DeepPep as core models, was proposed to obtain higher affinity binding peptide for a given target. Our method offers promise for the efficient generation and acquisition of *de novo* peptides with specific activity, and from which potential *de novo* active peptides against specific targets can be obtained through subsequent screening pipelines.

## Method

The overall design scheme is shown in Figure 1. It mainly includes the following three steps: 1) firstly train the peptide generation model through the general peptide dataset; 2) finetune the training with known active peptides to obtain a specific active peptide generation model and generate a large number of specific active peptides; 3) with the help of peptides Segment 3D modeling, docking, deep learning protein-peptide prediction model (optional), molecular dynamics simulation and other technologies to screen, obtain new potential designated active peptides, and finally submit to experimental verification. It is also possible to iterative such a process to obtain higher affinity peptides for a given target.

**Figure 1.**
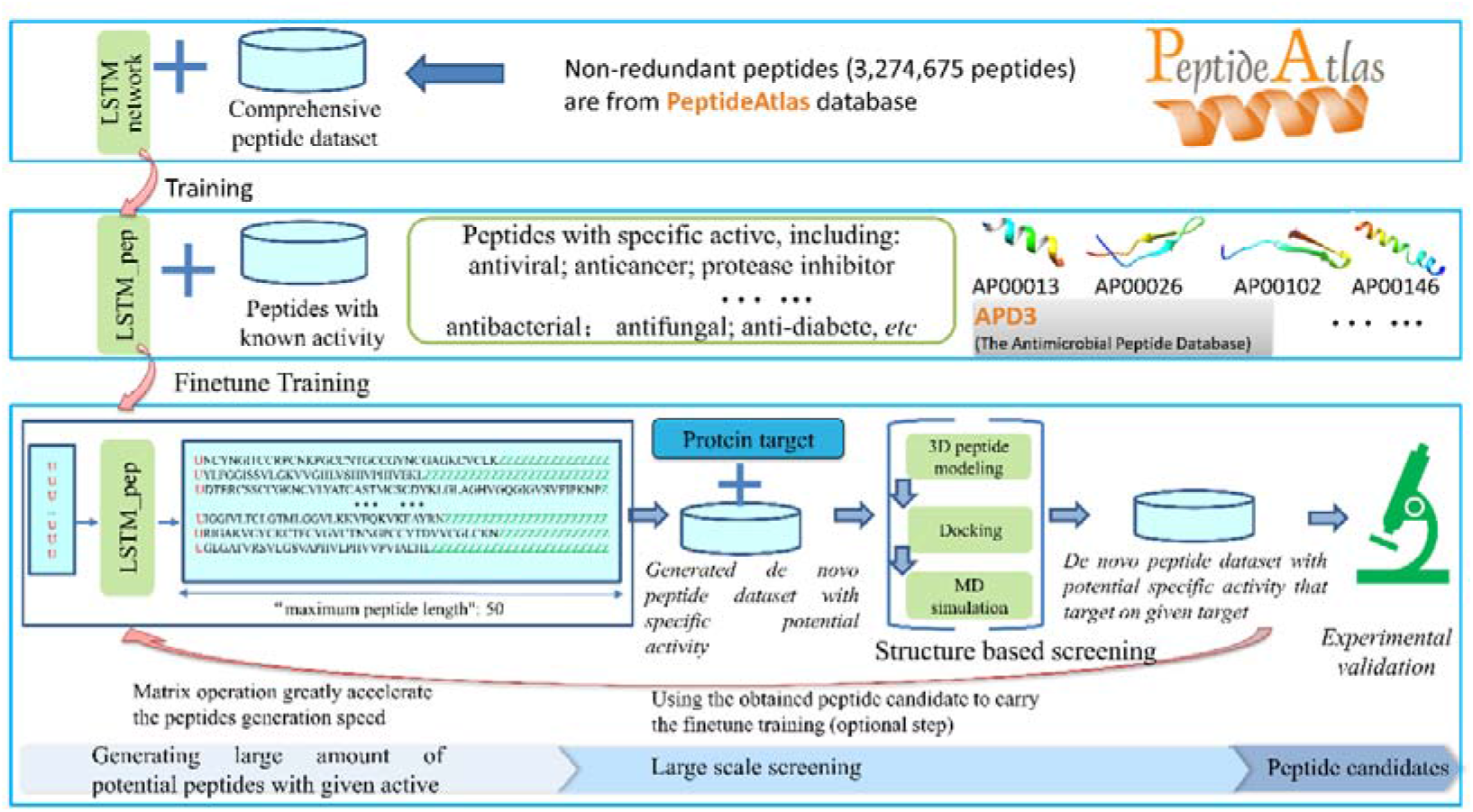
Overall flow chart. It mainly includes constructing general peptide generation models, finetuning specific active peptide generation models, and new peptide screening for specified targets.

### Collect peptide data for training

We obtained many peptides from the PeptideAtlas database[34] (http://www.peptideatlas.org/). By removing repetitive peptides and peptides containing non-standard amino acids, we finally obtained 3,274,675 peptides and used them as a training set to train a general peptide generation model. Then we convert the peptides into a matrix, and the amino acids in the matrix are represented in the form of one-hot. The peptide collection and preparation process are shown in Figure 2.

**Figure 2.**
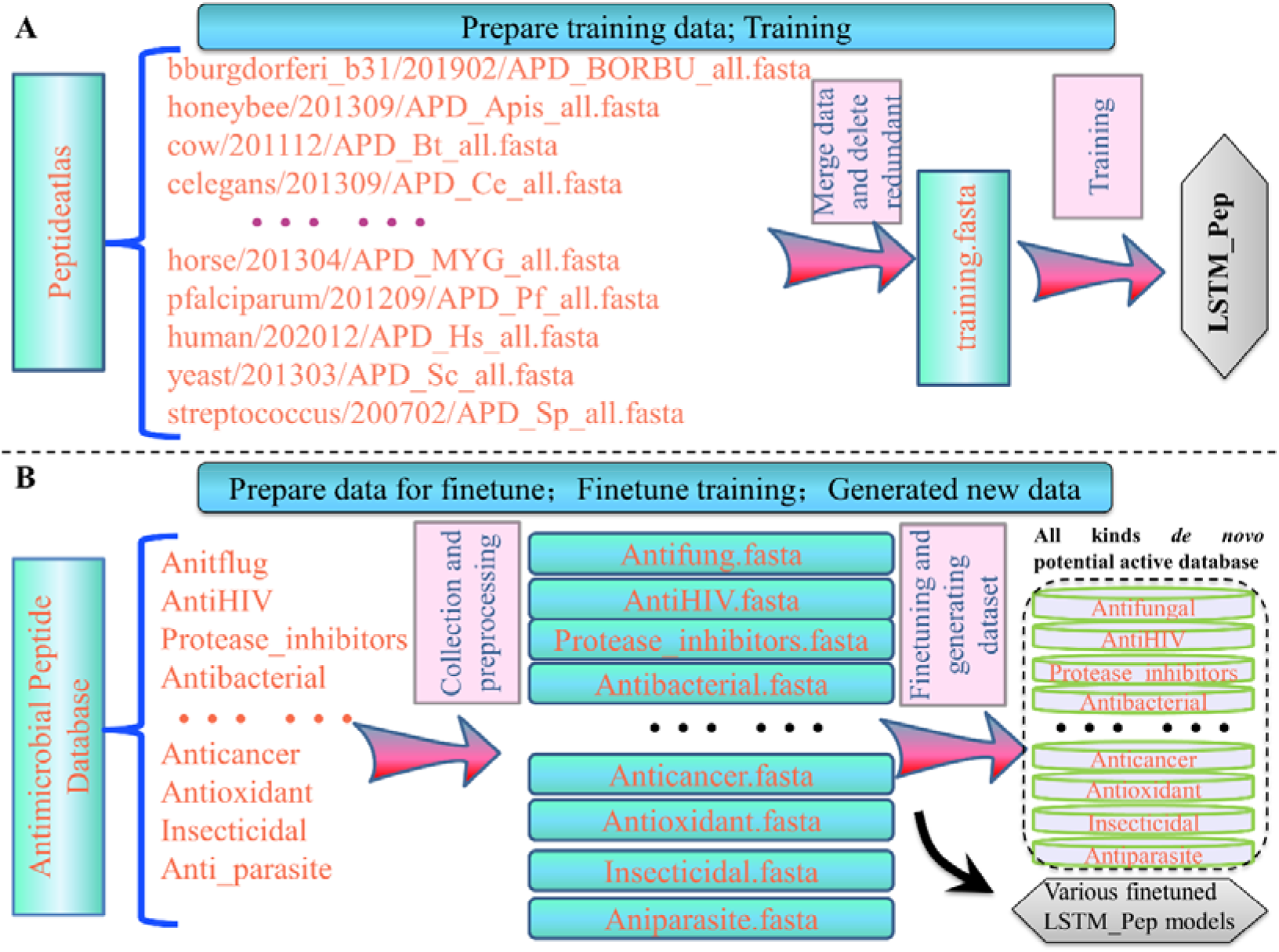
Shows the procedure for training and finetuning the LSTM_Pep model. A, shows the collection and preprocessing of training data; B, shows the preparation of data collection for finetuning training and generating a variety of *de novo* potentially active peptides.

### Peptide generation Model construction and training

The model structure used for training is a standard LSTM structure, as shown in Figure S1. The model consists of two layers of the LSTM network, one fully connected layer, and finally output through softmax. The first and second LSTM layer networks contain 256 nodes, and dropout is selected as 0.5 and 0.3, respectively. A multi-class loss function (categorical_crossentropy) is used in training. In the generic peptide model generation training, the epoch is set to 22. The model structure is adopted from the LSTM_Chem[18].

### Collecting peptide datasets of known biological activity and perform model retraining

We obtained a database of known active peptides from the Antimicrobial Peptide Database[1]. The database contains a variety of peptide datasets with different biological activities, including antifungal, anti-HIV virus, antibacterial, protease inhibitors, anticancer (antiviral), antiviral, insecticide, anti-diabetic, anti-parasite, *etc*. Of course, there are other active peptide datasets in the future, and finetuning training can also be performed according to this method.

During the migration training process using finetune, the structure of the training model is consistent with the original training institution. The starting model and weights are derived from the model trained in the first step, and the given active peptide data is used as training data for short-term finetuning training. (Epoch setting 12).

### Collection of generated novel peptides with specific potential activities

After the model was retrained for each selected biological activity, we immediately generated 5,000 whole peptides with the new model obtained (this value can be adjusted according to user needs, for example, adjusting to 50,000 may generate more peptide fragments). After removing duplicates and non-original (duplicated with training set peptides) peptides, we obtained new peptide datasets with different potential activities ranging from 4210 to 5000 data, as shown in Table 1. These novel peptides have the potential to have similar biological activities to the training sample peptides. Therefore, finding specific active peptides from these new peptide libraries is expected to reduce the experimental time significantly. However, the traditional method of finding the specified active peptides from random peptides will take a long time and many resources. Even if the peptides with a length of 8 need to be traversed, 20^8 experiments are required. In short, this kind of new potential peptide database will greatly contribute to the discovery of active peptides in further experiments in the future.

**Table 1.** The number of active peptides used for finetuning generated unique potential active peptides and the number of dissimilar peptides after removing 60% similar peptides.

| **Biological activity** | **The number of**<br>**corresponding**<br>**active peptides**<br>**used for**<br>**finetuning**<br>**training** | **The number of**<br>**completely new**<br>**active peptides**<br>**after**<br>**de-redundancy** | **The number of**<br>**dissimilar after**<br>**removing 60%**<br>**of similar**<br>**peptides** |
| --- | --- | --- | --- |
| Antifungal | 1210 | 4793 | 1697 |
| Antibacterial | 2742 | 4825 | 1533 |
| Antibiofilm | 66 | 4917 | 1034 |
| Anticancer | 253 | 4425 | 672 |
| Anti-diabetic | 16 | 4997 | 1244 |
| AntiHIV | 109 | 4852 | 1375 |
| Antimalarial | 33 | 5000 | 4302 |
| Anti-MRSA | 183 | 4893 | 1354 |
| Antioxidant | 28 | 4740 | 1261 |
| Anti_parasite | 137 | 4942 | 1671 |
| Anti_TB | 14 | 4994 | 2124 |
| Anti-toxin | 15 | 4201 | 645 |
| Antiviral | 193 | 4848 | 2003 |
| Insecticidal | 40 | 4821 | 808 |
| Ion_channel | 7 | 5000 | 4456 |
| Protease_inhibitors | 31 | 4998 | 3561 |
| spermicidal | 14 | 5000 | 2992 |
| surface_immobilized | 31 | 4980 | 922 |
| Wound_healing | 23 | 4970 | 1837 |

### Construction and training DeepPep Model for predicting Protein-peptide interaction

The protein-peptide positive data was from the PepBDB dataset. The negative data was constructed by assuming that randomly selected peptide is often not binding with a given target. We cross-docked the protein with randomly selected three peptides, and for each docking, we kept three conformations. All the docked conformation is taken as the negative dataset. Notably, we replicated the positive data 9 times as an oversampling scheme. Finally, we obtained 151,603 training data (positive 8500*9, negative 75103), 10,694 validation data (positive 600*9, negative 5,294), and 6,400 testing data (positive 712, negative 5,688). Since the negative data are much more than the positive data. To keep the balance between the positive and negative data, we have replicated nine times of positive data in the training and validation set. We extracted protein-peptide residue contact pairs with cutoff of Cαdistance 1nm. To capture spatial information for interacting pairs, we used K-Means methodology-derived spatial coordination in scikit-learn software to cluster protein residue into five groups; too few groups will lead to excessive loss of spatial information, whereas too many groups will lead to very few atoms within one group[35]. During the input file preparation step, residue pairs belonging to the same group were kept near one another to maintain some neighbor information of the protein residues. The one-hot representation of each residue type within residue pairs was concatenated on the same line. The concatenated representation of the pairs was written into files line by line. The maximum line number was 300 to provide coverage for most of the pairs. To standardize the input format, for any pair number less than 300, lines were padding with zeroes, and for any pair number larger than 300 (243/8500, 2.86%), the post-300 parts were omitted. In such way, we obtained figure-like matrix as input data. Then the model was carried out training, validation, and test.

### Showcase the application of obtained peptide dataset in *de novo* peptides screening by traditional docking and MD simulation

After obtaining new peptides, we still need to spend a lot of time verifying these peptides’ activity using experimental methods. It is still too resource-intensive for many small laboratories. The final target information is still completely unknown. We select antiviral peptides as proof-of-concept cases for screening and gradually narrow the range of peptides targeting specific targets through computer-aided screening methods. To facilitate the display of the screening process, we selected SARS-COV-2 main protease[36], an anti-COVID-19 therapeutic target, as an example, and screened new antiviral peptides against this target from the generated new peptide library, as shown in Figure S2.

The main protease is obtained from PDB data (PDB ID 6Y2F[37]). Various methods such as modeling, traditional docking, deep learning, and pocket molecular dynamics (MD) simulation are gradually used in the screening process to achieve high efficiency and complementary advantages. The trRosetta modeling[33], ZDOCK docking and rosetta docking procedure are shown described in the section 1 of the Supplementary material. The detailed pocket MD simulation and metadynamics simulation procedure are described in the section 2 of the Supplementary material, which is similar to our previous work[30].

## Result

We obtained generated peptides for each bioactive. The distribution of the number of different potential active peptides generated is shown in Figure S3. We can see all the 18 peptides have generated more than 4000 unique peptides. We find the Anti-diabetic, Antimalarial, Ion_channel inhibitor, and spermicidal obtained 5000 potential peptides. Among the generated potential designated active peptides, we obtained 4848 potential antiviral peptides, 4425 anticancer peptides, and many other types of active peptides. We also examined the peptide length distribution, and among all the obtained peptides, different lengths were shown in Figure S4. The average peptide length ranged from 15 to 40, corresponding to different types.

To examine the diversity of generated peptides, we clustered peptides with different potential activities. After removing peptides with a similarity greater than 60%, we found that most datasets still retain many samples, as shown in Table 1.

Taking new potential antiviral peptides as an example, we built their 3D structures, such as some potential antiviral peptides, as shown in Figure S5. We can see the diversity of these structures.

### The predicted peptides for the main protease

Screening of peptides targeting specific targets from the obtained new peptide library can narrow the scope of subsequent experimental verification and clarify the particular mechanism of action. The specific calculation process includes peptide modeling, target protein pocket positioning, ZDOCK rigid docking, rosetta flexible docking, and molecular dynamics simulation.

After ZDOCK rigid docking, and rosetta flexible docking, we obtain 22 compounds with rosetta total_score <=-454.8 kJ/mol. The 22 protein-peptide complexes were carried out in a pocket MD simulation. Among them, 8 peptides (antiviral_430, antiviral_490, antiviral_822, antiviral_1443, antiviral_1712, antiviral_1996, antiviral_3445, antiviral_4465) that have average RMSD value <= 0.75nm during last 20ns simulation was selected and shown in the Figure S6A. The number of hydrogen bonds was also shown in Figure S6B; most of them have several stable hydrogens formed during the MD simulation.

We also calculated the free energy landscape for the 22 protein-peptide complexes, shown in Figure S7. We find that six peptides (antiviral_1947, antiviral_2280, antiviral_3258, antiviral_430, antiviral_4616, antiviral_88) are not favored to bind on main protease according to the free energy landscape. The peptide antiviral_3445 also has no clear tendency to prefer binding according to the free energy landscape. However, most other peptides prefer to bind to the Main protease.

We obtained six peptide candidates (antiviral_490, antiviral_822, antiviral_1443, antiviral_1712, antiviral_1996, antiviral_4465) for the Main protease that fulfilled low RMSD during pocket MD simulation and have preferred binding according to the free energy landscape. We analyze the 3D pocket of the Main protease with those peptides from the last frame of the 100ns MD simulation, shown in Figure S8. There are many close contact residues pair (residues distance within 0.3nm), established in Table S1.

The flexible docking method, such as rosetta, is not efficient enough, especially when we want to iterative generating and screening. Furthermore, its score function has a big approximation, which may lead to not accuracy in many situations. To solve this problem, we developed a Deep learning-based binary model for protein-ligand interaction prediction, shown in Figure 3. The model performance in training, validation and test set is listed in the Table 2, we observed that the model has AUC value of 0.8673, TPR value of 0.8652, and MCC value of 0.3239 in testing sets, indicating the model have well performance in identifying native-like protein-peptide complex. Notably, the data in the testing set have very unbalance distribution, which the negative data set are much large than positive dataset. This is closer to the real application scenario that unbinding peptides usually are much easy encountered during screening. After we obtain the model, we can use it to do the screening; noticeably, our method depends on ZDOCK to generate conformations; since dock score may not be accurate enough, we need to predict more top score conformations and select one top prediction as the final protein-peptide binding possibility score, shown in Figure 3C.

**Table 2.** The performance of DeepPep on training, validation and test data set.

| Data | AUC | Accuracy | TPR | Precision | MCC | Pos_num | Neg_num |
| --- | --- | --- | --- | --- | --- | --- | --- |
| Training | 0.9799 | 0.8413 | 0.9995 | 0.761 | 0.7188 | 8500*9 | 75103 |
| Validation | 0.8581 | 0.7488 | 0.8633 | 0.7053 | 0.5098 | 600*9 | 5294 |
| Test | 0.8673 | 0.667 | 0.8652 | 0.2324 | 0.3239 | 712 | 5688 |

**Figure 3.**
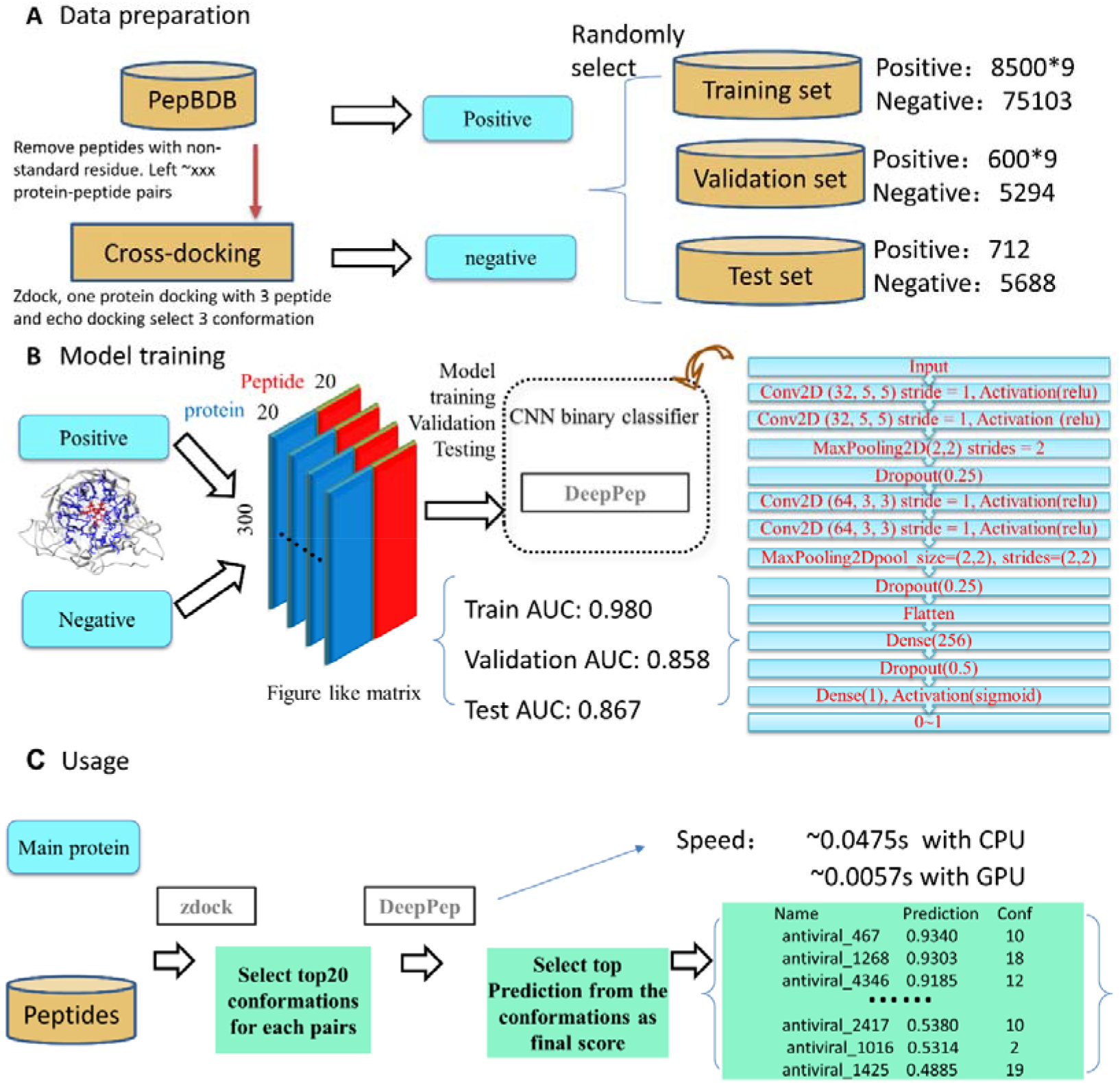
The procedure of constructing the protein-peptide model and its usage in screening. A, data preparation; B, model training, validation, and testing; C, Example of model usage.

## Discussion

An LSTM model was used for peptide generation, and an iterative generative screening method was used to efficiently obtain active peptides for a given protein. The present work utilizes the peptide database to train a stable peptide generation model. After that, the Antimicrobial Peptide Database data were used for finetuning learning to obtain a special model for generating designated active peptides. In addition, the work points out the subsequent use of the generated new active peptide library for screening against specific targets. A possible iterative generation screening scheme is also pointed out, which provides a process and ideas for designing ultra-high-affinity active peptides. A rapid protein-peptide binding prediction model is also our new method to improve the accuracy and efficiency of screening.

### Iterative generating and screening

Considering that one round of generating and screening peptides against a given target sometimes might make it hard to achieve the desired strong binding affinity. We try to iterative generating and screening, which the input data for finetuning are from previous round selected candidates. In such a way, it is possible to obtain higher and higher affinity candidates. However, such iteration would be more practical if we used faster screening tools; we chose the newly developed DeepPep as the final screening tool. Take the SARS-CoV-2 main protease as an example, we iterative generating peptides by integrating LSTM_Pep, tr_rosetta modeling, Zdcok docking, and DeepPep screening, as shown in Figure 4. Furthermore, we also selected some high DeepPep score main protease-peptide complexes for MD simulation and Metedyanmics simulation to further examine their interaction details, binding stability and binding free energy landscape, shown in Figures S9,10,11,12.

**Figure 4.**
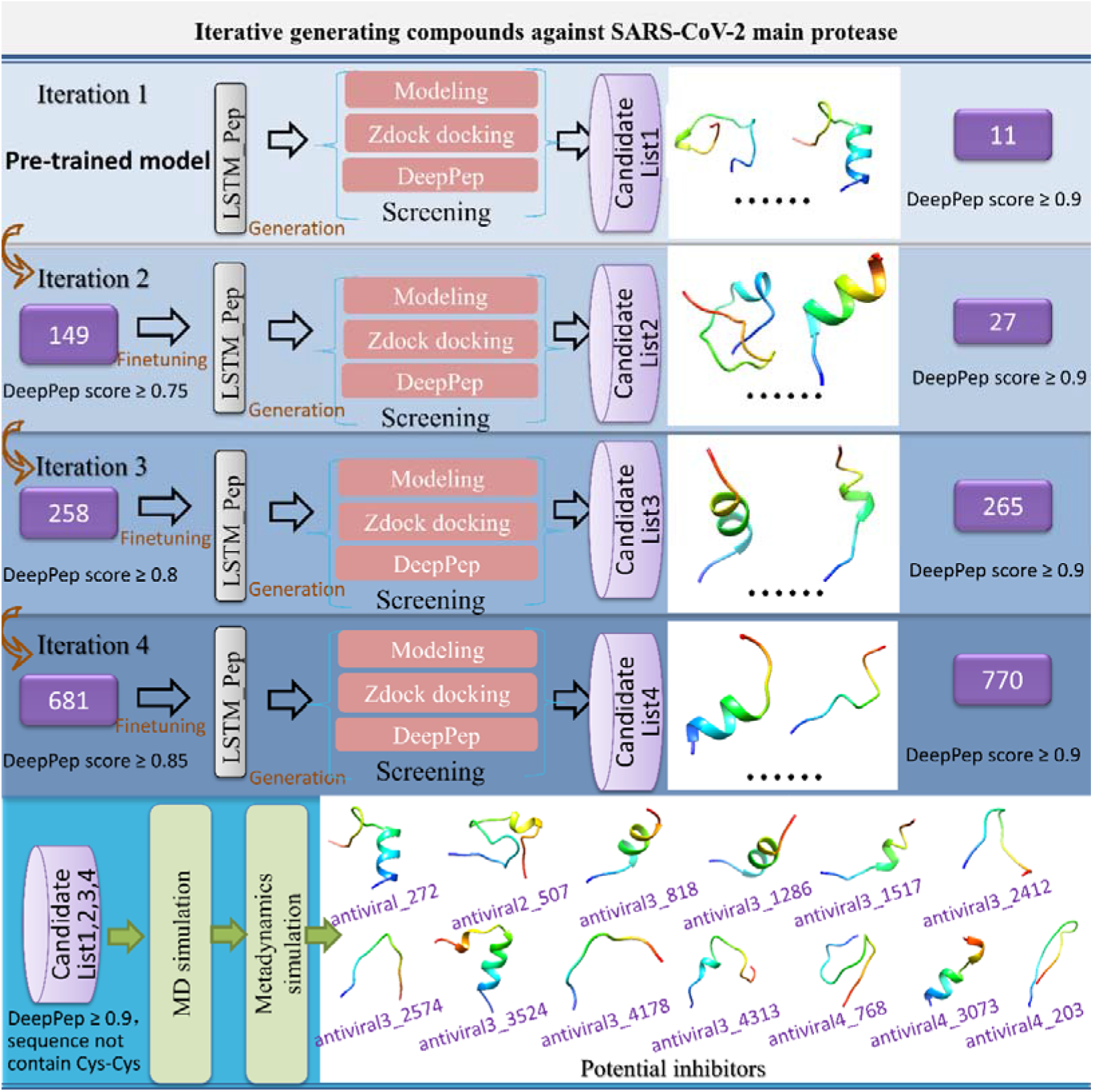
Iterative peptide generation and screening for a given target. For each iteration, the previous round high potential binding candidates are used as finetuning data, in such a way, more and more high potential binding candidates are kept. The candidates are finally selected to carry out MD simulation and metadynamics simulation to check the binding stability, interaction details, and free binding energy landscape.

### The generated bioactive peptides can be widely used in an application

With this our proposed strategy, we can screen various peptides with different bioactive against various protein targets. Take antiviral peptides, antibacterial, and anticancer peptides discovering as an example, we can obtain lots of *de novo* peptides against many different targets from many different kinds of viruses, such as RNA virus[38] (*e*.*g*. Coronavirus, HIV, HCV, *etc*.) and DNA virus[39] (*e*.*g*. herpes zoster, HBC, adenovirus, *etc*.). Take Coronavirus[40] as an example, the targets can be RNA-dependent RNA polymerase (RdRp)[41], Main protease, Nsp13[42], *etc*., as shown in Figure 5A. Also, take antibacterial as an example, we can obtain *de novo* peptides for different kinds of bacteria including gram-positive bacteria[43] (Staphylococcus aureus, Streptococcus pyogenes, and Strep. Pneumoniae, etc.) and gram-negative bacteria[44] (Klebsiella, Pseudomonas aeruginosa, Acinetobacter, Escherichiacoli), as shown in Figure 5B. Take the anticancer as an example, we can obtain *de novo* peptides for breast cancer[45], lung cancer[46], prostate cancer[47], *etc*., and the targets can be BCL-2[48], TOP1[49], CDK4/6[50], *etc*., as shown in Figure 5C.

**Figure 5.**
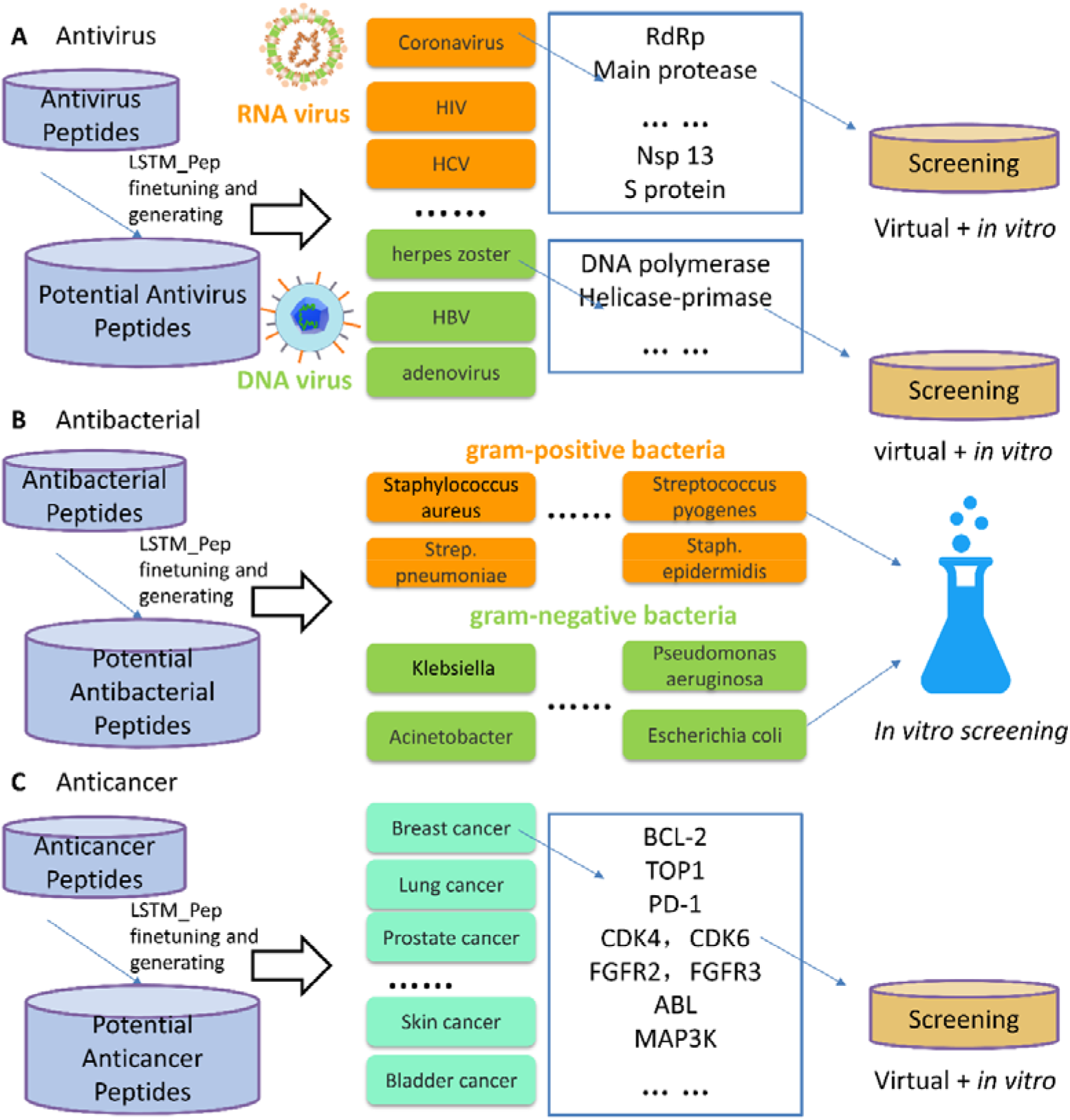
The huge potential of generated *de novo* peptides in developing various kinds of bioactive peptides. A, the potential usage of our methods in antivirus; B, the potential usage of our methods in antibacterial; C, the potential usage of our methods in anticancer.

## Conclusion

Inspired by small molecule generation software such as LSTM_CHEM, we developed a peptide generation model. At the same time, we used the known active peptide deduplication training model and generated a large number of brand-new potential active peptides through the obtained retrained model. As we know, compared with compounds, peptides are easier to synthesize and cost less to acquire. Therefore, this kind of calculation method will be of great value to the development of peptides. We also propose a pipeline for screening these novel potential active peptide libraries for specified protein targets, which can be used to narrow the scope of subsequent peptide research further. This directly points to a novel approach to efficiently obtain peptides targeting specified targets. The screening pipeline plays at least two roles.

On the one hand, it dramatically reduces the scope of the candidate list for experiments and improves accuracy. The other makes us understand the underlying mechanism of how the peptide shows bioactive, which would be important for later structure-based peptide modification. Finally, the iterative generation screening scheme can be used to optimize peptides further; that is, using the peptides screened in this iteration as the input for the next finetuning training, it is expected to obtain high-affinity active peptides targeting the specified target.

## Key Points

- We have developed a deep learning-based model that can generate native-like peptides. Through mode finetuning, we can generate various kinds of potential *de novo* bioactive peptides, including Antifungal, Antibacterial, Anticancer, Anti-diabetic, AntiHIV, Antimalarial, Antioxidant, Anti_parasite, Anti-toxin, Antiviral, Protease_inhibitors, *etc*. This can greatly promote de novo bioactive peptide discovery. Furthermore, we have proposed a hybrid pipeline to further obtain potential bioactive peptides against a given target, which contains deep learning, docking, MD simulation and Metadynamics simulation. The known protein-peptide structure interaction details also help late-stage peptide design and refinement.
- We have developed a deep learning protein-peptide prediction model that can be used to efficiently and accurately screen generated peptides for a given target. Moreover, the present work demonstrates that deep learning-based iterative generation and screening produce higher predicted affinity *de novo* peptides candidates for a given protein target. The iterative peptide generation and screening strategy proposed here would be highly desirable for pharmacological companies to efficiently obtain potential de novo peptide drugs that can apply both structure and functional patents.
- Using the Main protease as a proof-of-concept example, we have obtained several peptide candidates. The predicted candidates were further examined by predicted binding stability and predicted binding free energy landscape from molecular dynamics simulation and metadynamics simulation. Moreover, we find combined LSTM_Pep and DeepPep can generate and screen *de novo* active peptide very efficiently, and applied these two methods in target Xanthine oxidase led to successfully discover an active *de novo* peptide (ARG-ALA-PRO-GLU).

## Availability of data and materials

The proposed models and the scripts are available in GitHub public repositories (https://github.com/haiping1010/New_peptide_iteration). All other data, data preparing code, model source code, model application code that requires to reproduce the result are available from the corresponding author upon request.

## Author contributions

H.Z. designed the study. H.Z., K.M.S, Y.Y., Y.W., X.W., Y.P., Y.J. performed computations and data analyses. All authors contributed to writing the manuscript. H.Z. and J.Z.H.Z. supervised the study. All authors read and approved the final manuscript.

## Acknowledgments

This study was supported in part by the National Science Foundation of China (Grant No. 62106253, 21933010), Shenzhen International Science and Technology Exchange and Cooperation Project (No. GJHZ20210705141803010) (XW), the Key-Area Research and Development Program of Guangdong Province (2019B020213001) (XW), Research Funding for Innovation Project of Universities in Guangdong Province (2018KTSCX192) (XW), the Shenzhen KQTD Project (No. KQTD20200820113106007), Research Funding of Shenzhen (JCYJ20200109114818703).

