## Supplemental file for "Deep-learning based bioactive therapeutic peptides generation and screening"

**Supplementary material section 1:**

**Detailed procedure of trRosetta modeling, zdock and rosetta docking**

The trRosetta was used to build 3 model of the *de novo* peptides. The procedure was following procedures: 1, using hhblits[1] to generate multiple sequence alignments (MSAs) file; 2, Predicting interresidue geometries from MSAs using a deep neural network; 3, Structure modeling from predicted interresidue geometries by pyrosetta[2].

The zdock was used to dock the peptides to the target protein. We define the docking pocket as the 1.2 nm within the predicted binding ligands, 1000 conformations were generated, and only the top prediction was used for next step rosetta flexible docking.

The rosetta was used to do the flexible docking, 5 conformation was generated for each docking, and the top score was kept. Finally, we chosen 22 top protein-peptide as top predictions for next step MD simulation.

The detailed script used for trRosetta modelling, zdock, and rosetta docking can be found in GitHub (https://github.com/haiping1010/New_peptide_iteration/tree/master/iteration_main_protease_Antiviral_pep).

**Supplementary material section 2:**

**Detailed procedure of pocket molecular dynamics and metadynamics simulation**

The initial protein-peptide complexes were from the top score conformation rosetta docking (or the top DeepPep prediction of Zdock docking conformation).

We use a pocket molecular dynamics simulation (pocket MD, Supplementary Fig. 11**b**) to facilitate the simulation process by only keeping the binding pocket region for simulation (the pocket was defined as residues within 1nm of the peptide). Binding free energy calculation can be estimated by metadynamics simulations to explore whether protein-peptide will bind in solution. Metadynamics relies on addition of a bias potential to sample the free energy landscape along a specific collective variable of interest [3,4]. Note that the binding free energy calculations from Metadynamics may only be suitable for detect the general trend of binding in virtual screening.

The pocket MD is same as the classical MD simulation, except that we only using the pocket region to reduce system size for simulation [5], which is inspired by a previous dynamic undocking (DUck) method [6]. An in-house script was used to extract the pocket region of the protein (here, we used 1.2nm within the binding peptide), the N terminal and C terminal ends were capped with the ACE and NHE terminals, respectively. We applied position restrains to the ACE and NHE terminals to maintain the relative conformation of the pocket. MD simulation was carried out by Gromacs with AMBER-99SB force field [7,8]. Firstly, we created a dodecahedron box and put the target-peptide complex at the center. A minimum distance from the protein to box edge was set to 1 nm. We filled the dodecahedron box with TIP3P water molecules [9], the counter ions were added to neutralize the total charge using the Gromacs program tool [10]. The long-range electrostatic interactions under the periodic boundary conditions was calculated with Particle Mesh Ewald approach [11]. A cutoff of 14 Å was used for van der Waals non-bonded interactions. Covalent bonds involving hydrogen atoms were constrained by applying the LINCS algorithm [12].

We performed the energy minimization steps with a step-size of 0.001ns, 40 ps simulation with isothermal-isovolumetric ensemble (NVT), and 10ns simulation with isothermal-isobaric ensemble (NPT) for water equilibrium. After that, a 100ns NPT production run (step size 2 fs) was carried out. The Parrinello-Rahman barostat and the modified Berendsen thermostat were used for simulation with a fixed temperature of 308 K and a pressure of 1 atm. RMSD and hydrogen bond number of the trajectory were calculated using Gromacs tools.

The simulation was continued using the metadynamics approach for exploring the free energy landscape. The interface coordination number of atoms of protein peptide complex was used as collective variable (CV). The protein-peptide interface coordination numbers correlate with the numbers of atom contact, and larger coordination number usually indicates that protein-peptide is in binding state.

The coordination number $C$ is defined as follows by Plumed:

$$C = \sum_{i \in A} \sum_{j \in B} S_{ij} \quad (1)$$ and

$$S_{ij} = \frac{1 - \left( \frac{r_{ij} - d_0}{r_0} \right)^n}{1 - \left( \frac{r_{ij} - d_0}{r_0} \right)^m} \quad (2)$$

In the simulation, $n$ was 6, $m$ was 12, $d_0$ was 0 nm and $r_0$ was 0.5 nm. $d_0$ is a parameter of the switching function. $r_{ij}$ is the distance between atom $i$ and atom $j$. The degrees of contacts between two groups of atoms can be estimated by above function(1) [13]. Metadynamics simulation for each protein-peptide system was performed for 40 ns. During the metadynamics simulation, Gaussian values were deposited every 1 ps with a height of 0.3 kJ/mol. The widths of the Gaussians were 5 for the coordination number. The free energy landscapes of the metadynamics simulations along the CV were generated by the Plumed program and plotted using Gnuplot [14].

### Supplementary Figures:

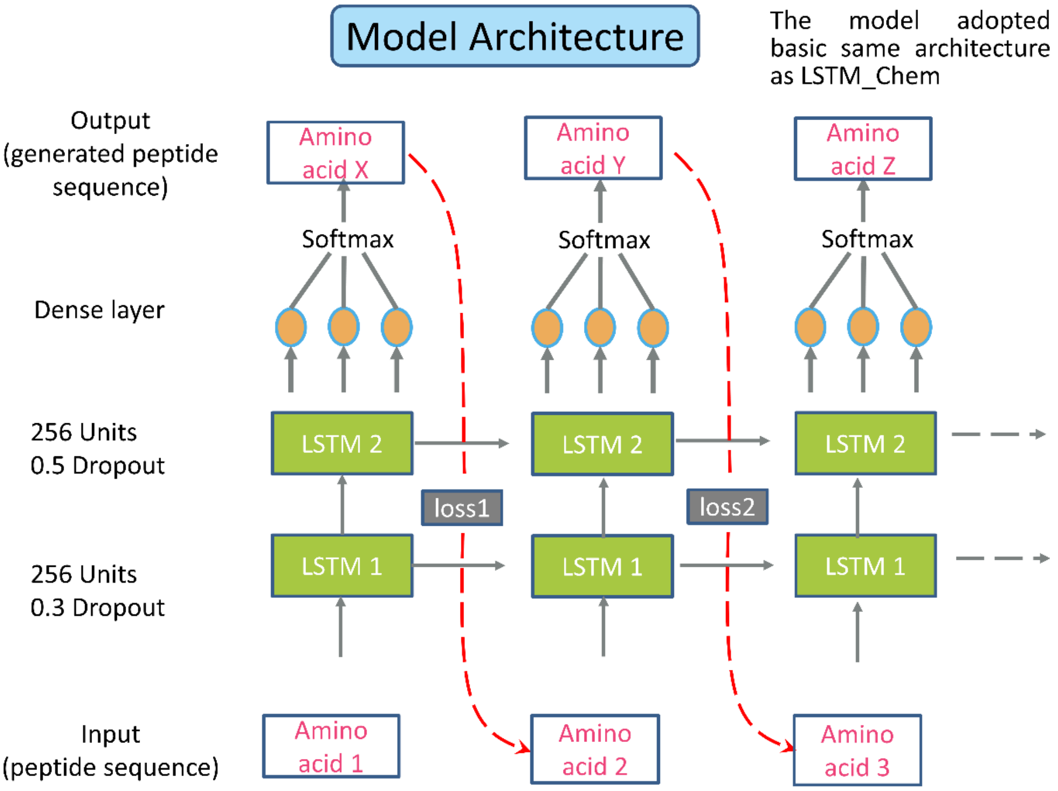


**Figure S1. The architecture diagram of the model LSTM_Pep.** The model adopts a structure consistent with the compound generation model LSTM_Chem.

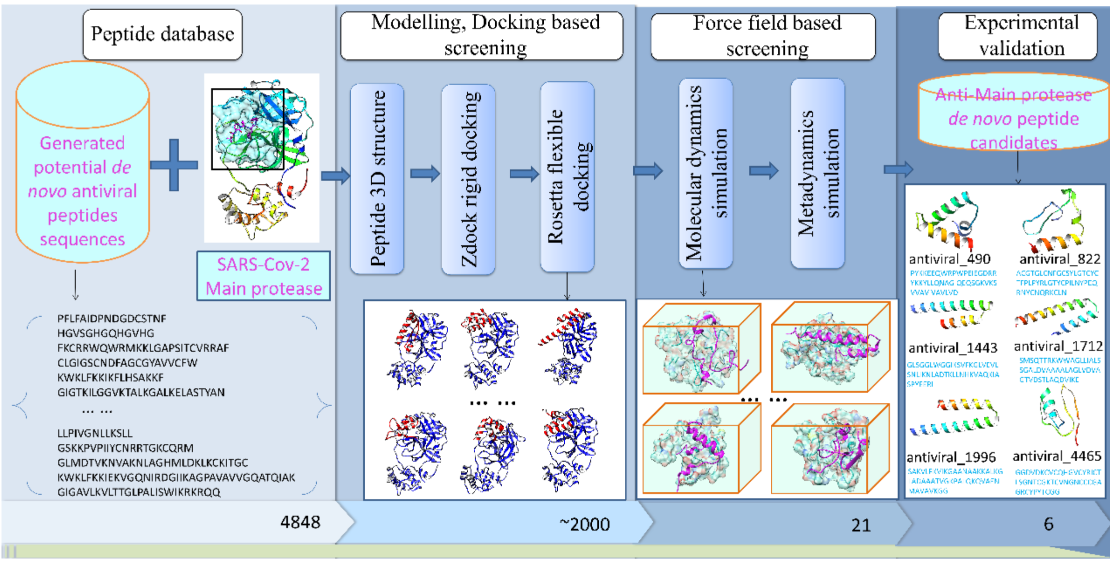

**Figure S2. Shows the procedure to further screen novel active peptides against specific targets from the generated potential active peptides.** The novel coronavirus therapeutic targets Main protease are used as examples.

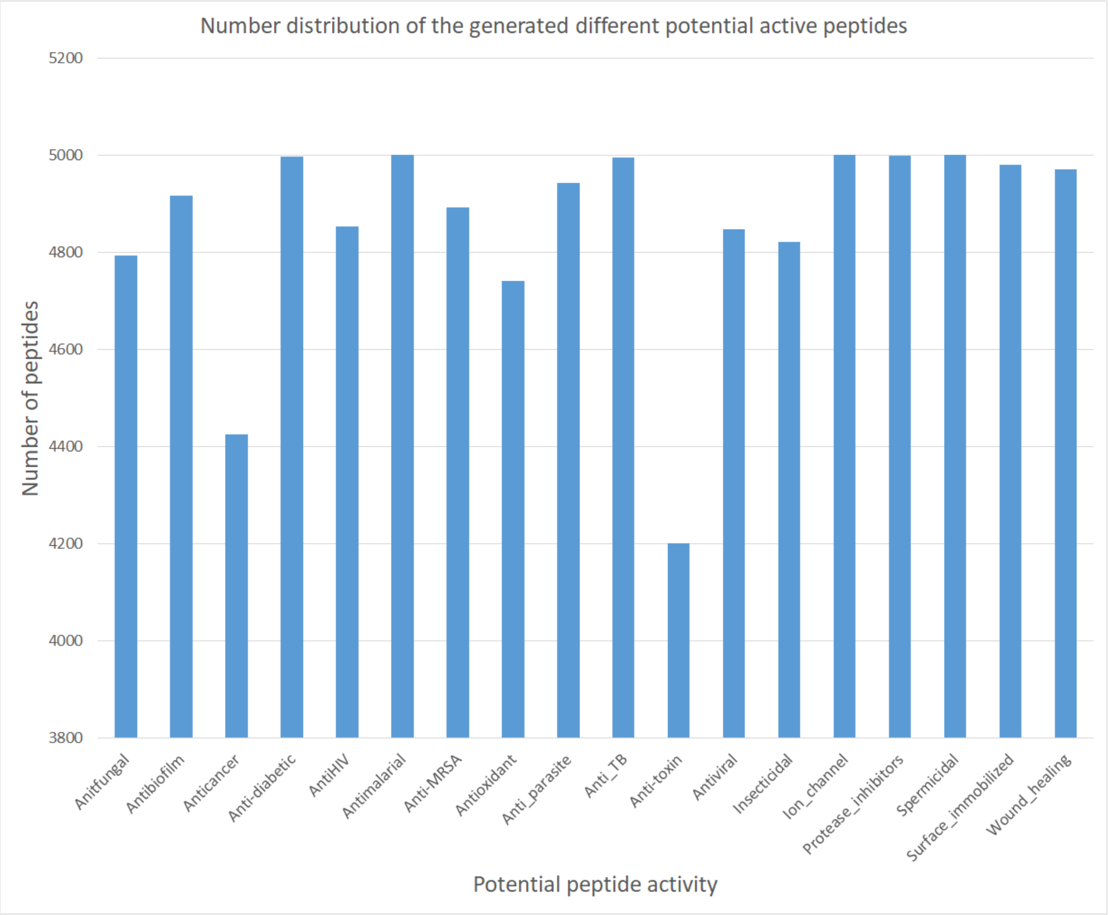


**Figure S3**. The number of different active peptides generated.

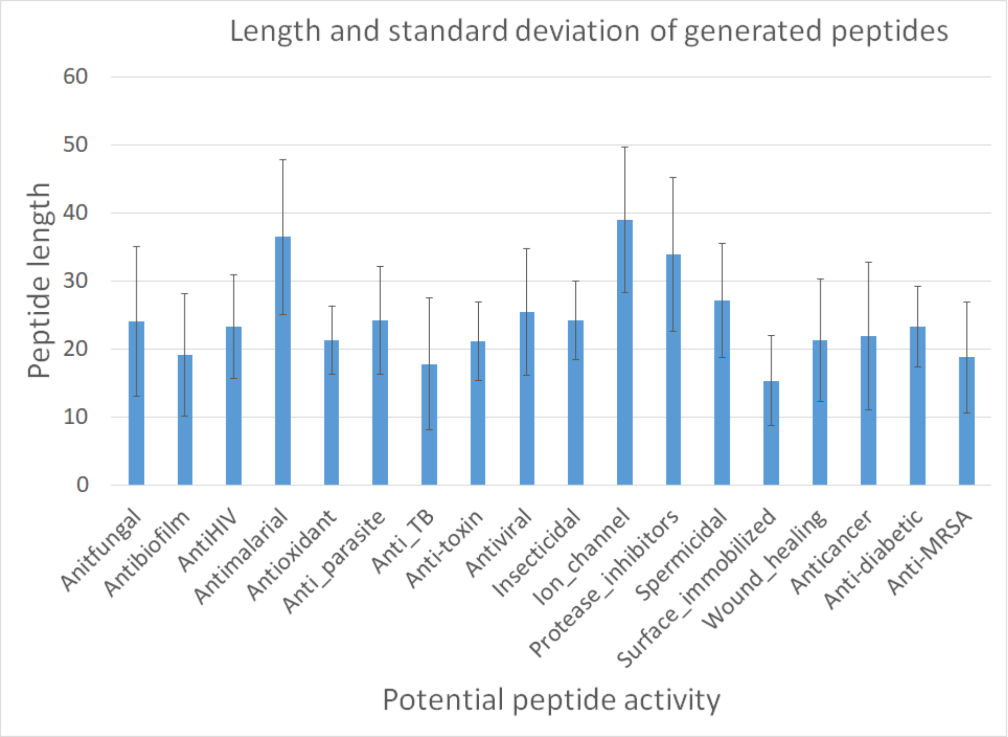


**Figure S4. The average length and standard deviation of various new active peptides were generated.**

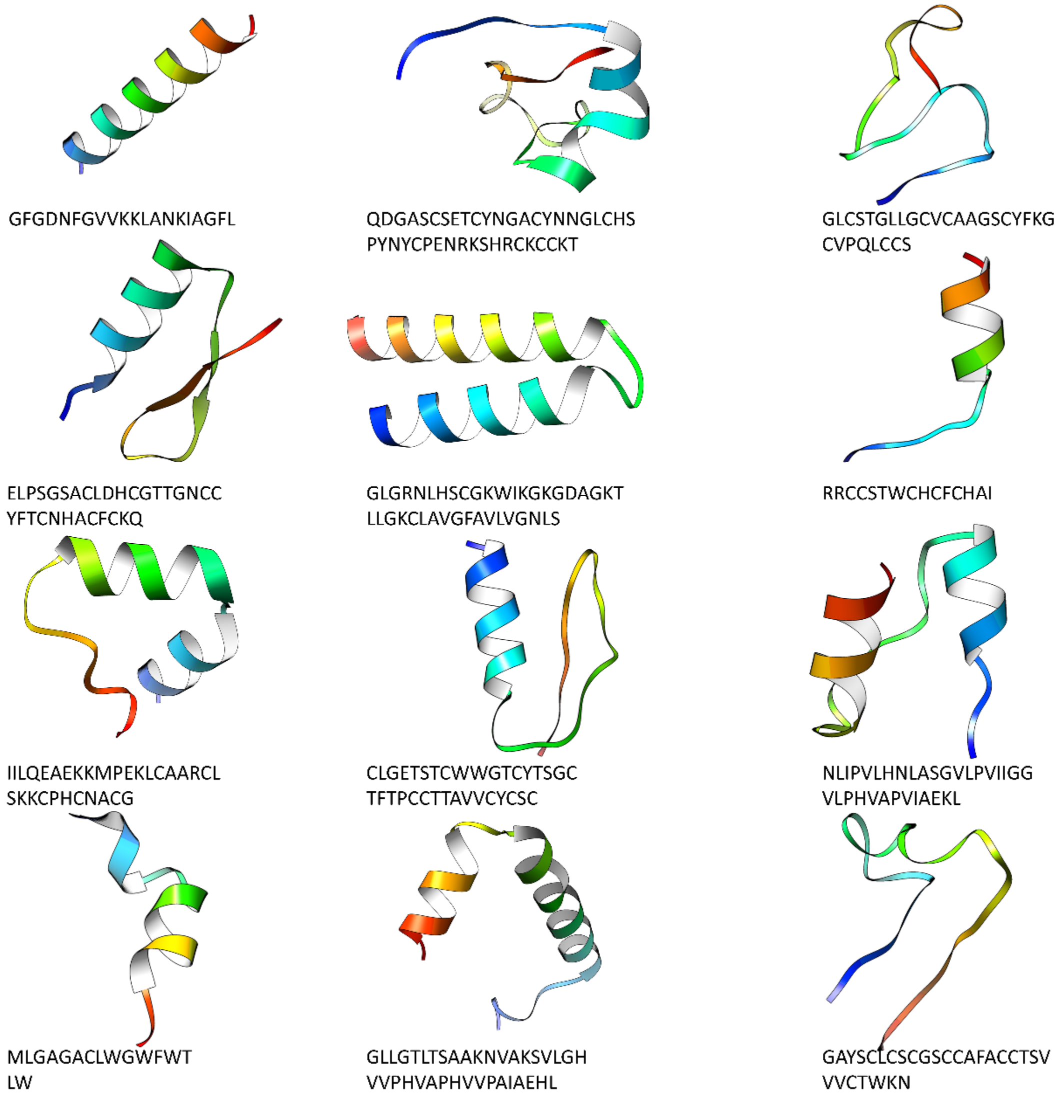


**Figure S5. Examples of potential antiviral peptides generated.**

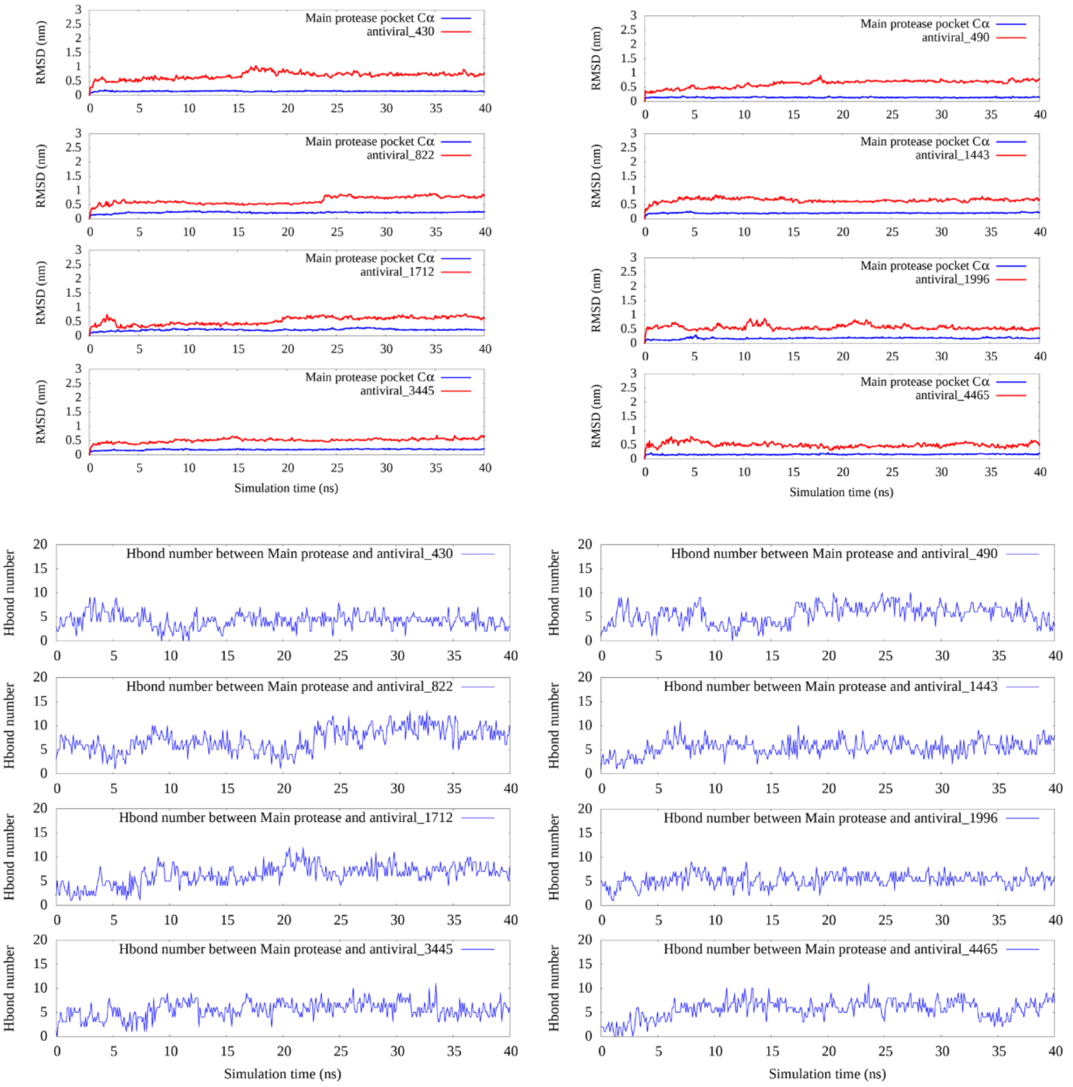


**Figure S6. The RMSD and hydrogen bind number for 8 selected protein-peptide complexes during MD simulation.** A. the RMSD of main protease pocket C alpha and the RMSD of the peptide during the pocket MD simulation. B. the hydrogen bond numbers between main protease pocket C alpha and the peptides during the pocket MD simulation.

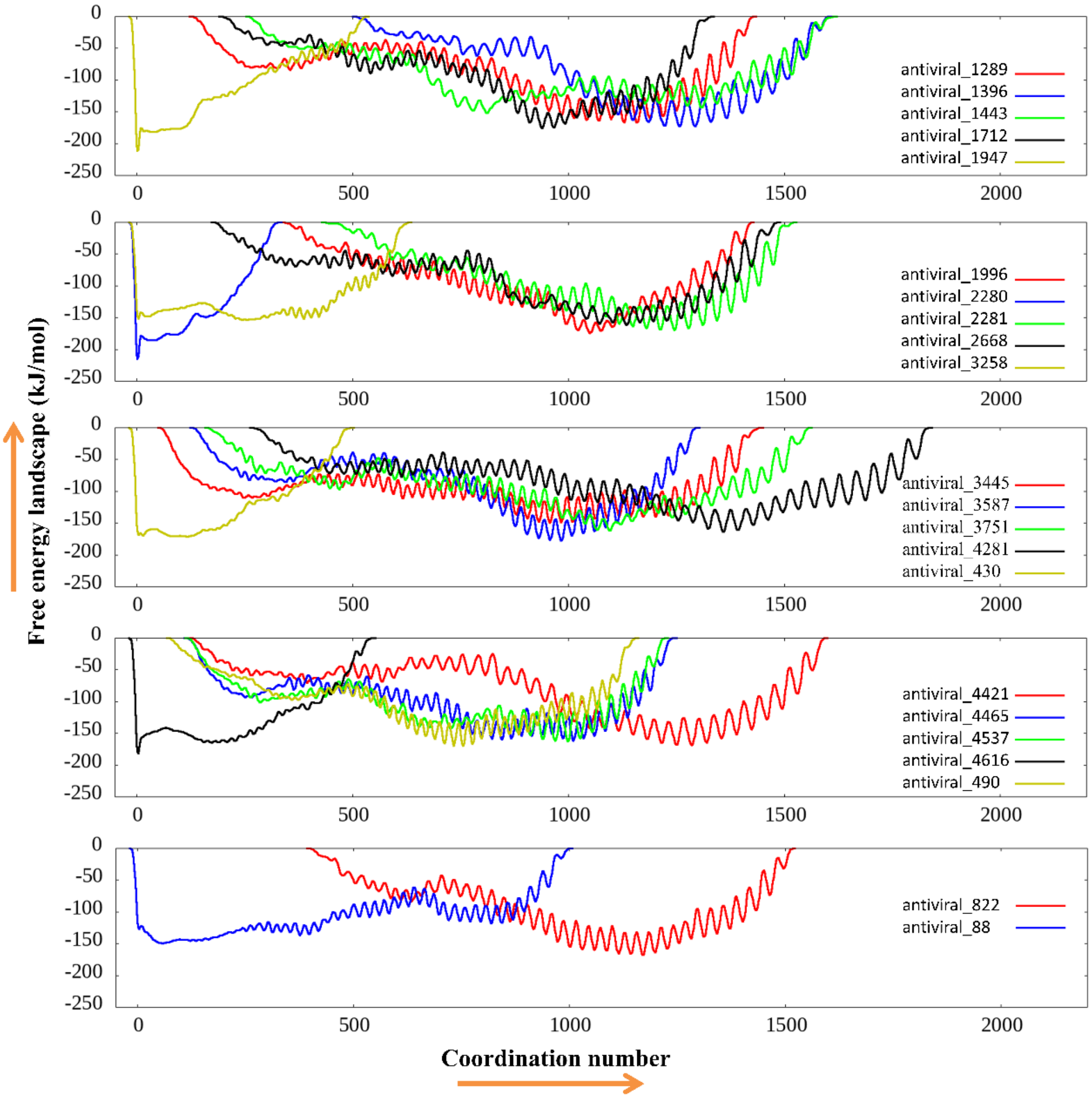


**Figure S7. The free energy landscape was calculated from the metadynamics simulation.**

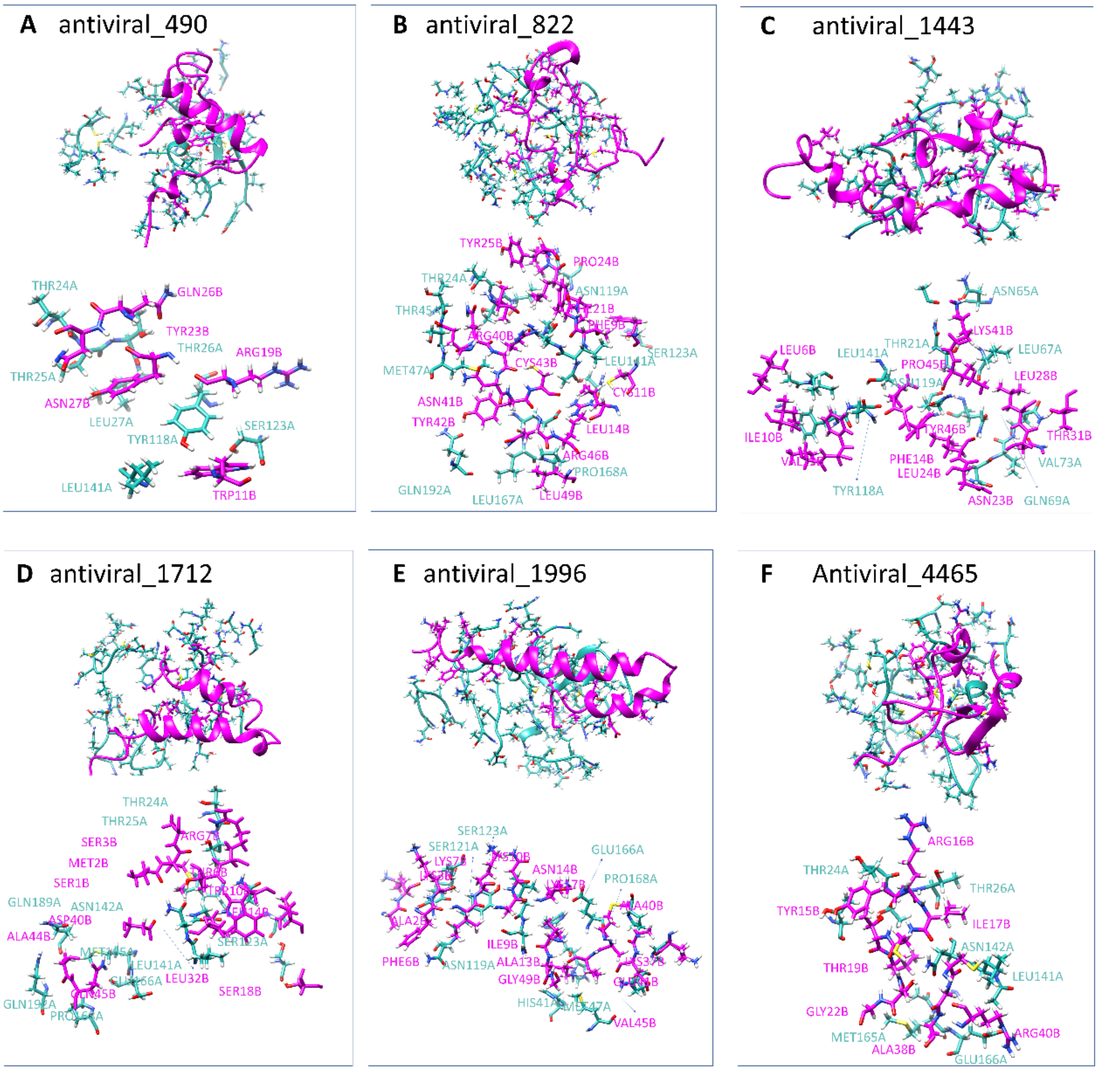


**Figure S8. The 3D conformation of the Main protease pocket with peptide from the 100 ns MD simulation.** The residue pairs within 2.5Å between protein and ligand were shown in atomic details, the protein was colored light blue, and the peptide was colored magenta.

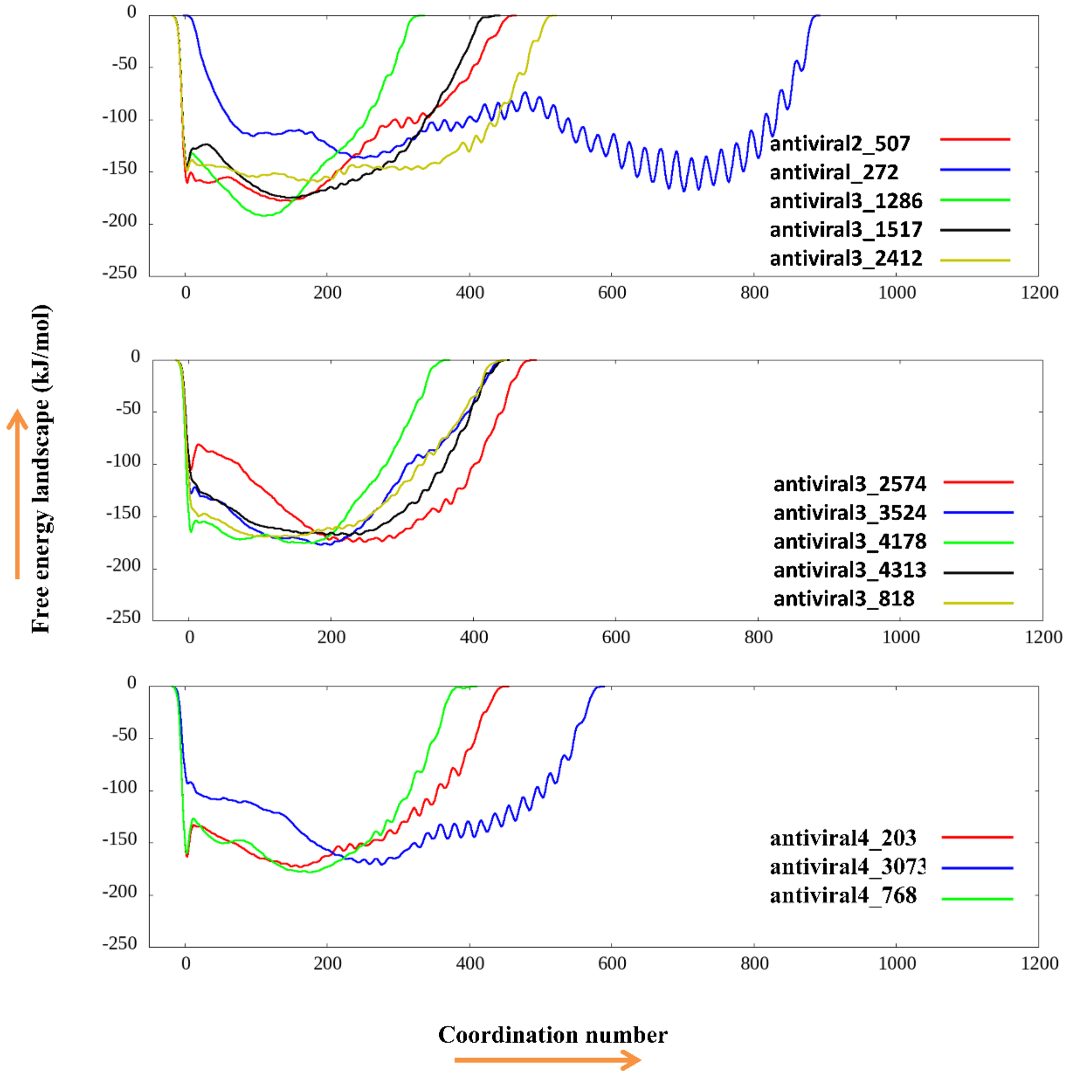


**Figure S9. Free energy landscape of 13 final selected peptides from Metadynamics simulation.**

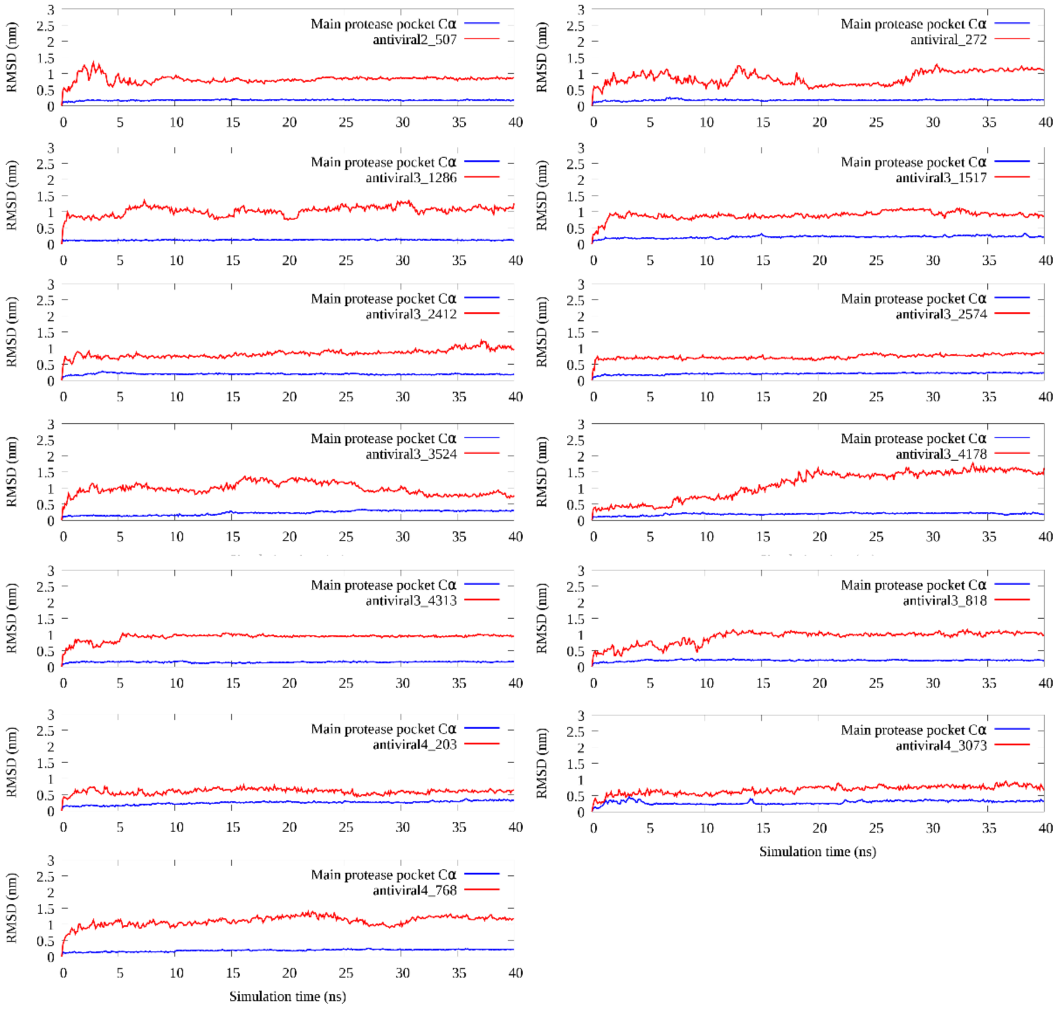


**Figure S10. The RMSD value of the 13 final selected peptides and their binding protein pocket from MD simulation.**

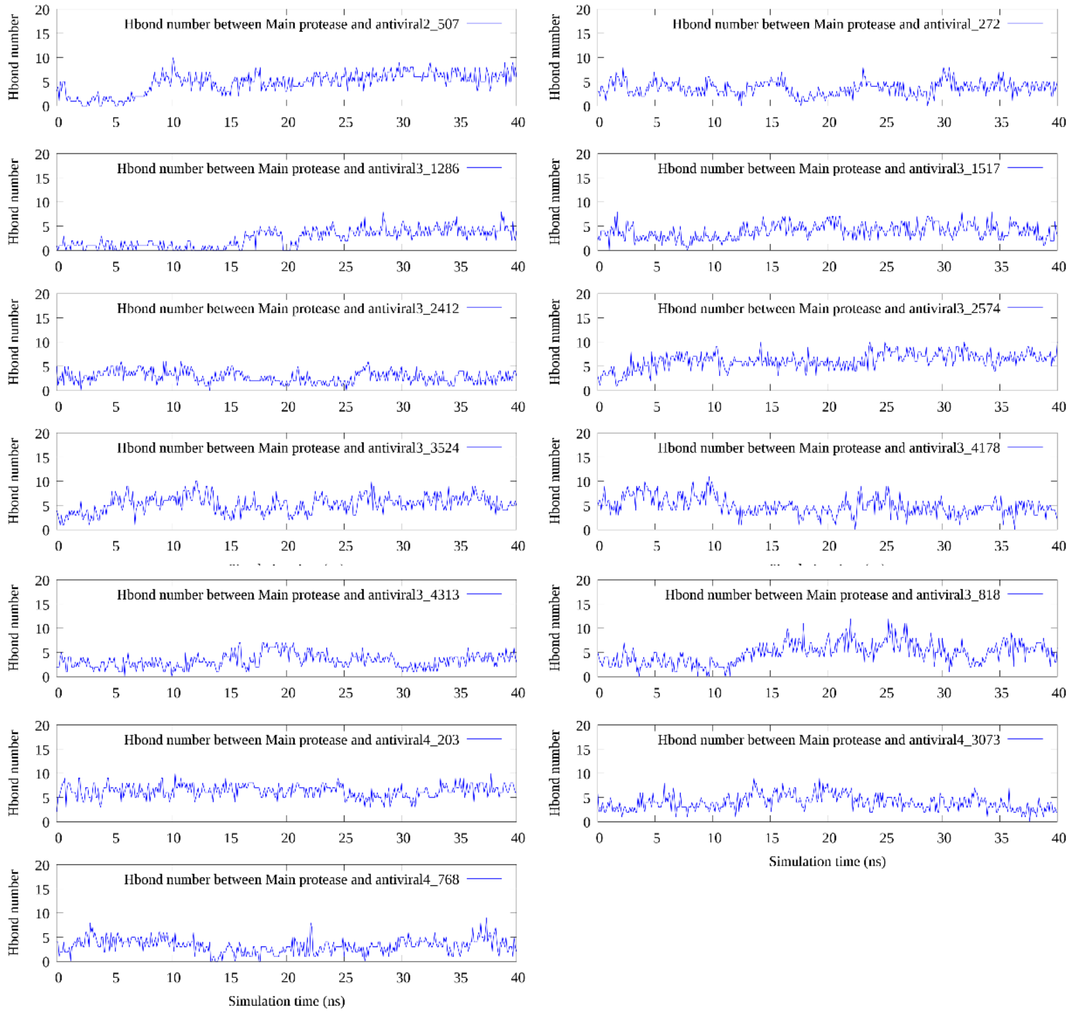


**Figure S11. The calculated hydrogen bond numbers of the 13 final selected peptides with their binding protein pocket from MD simulation.**

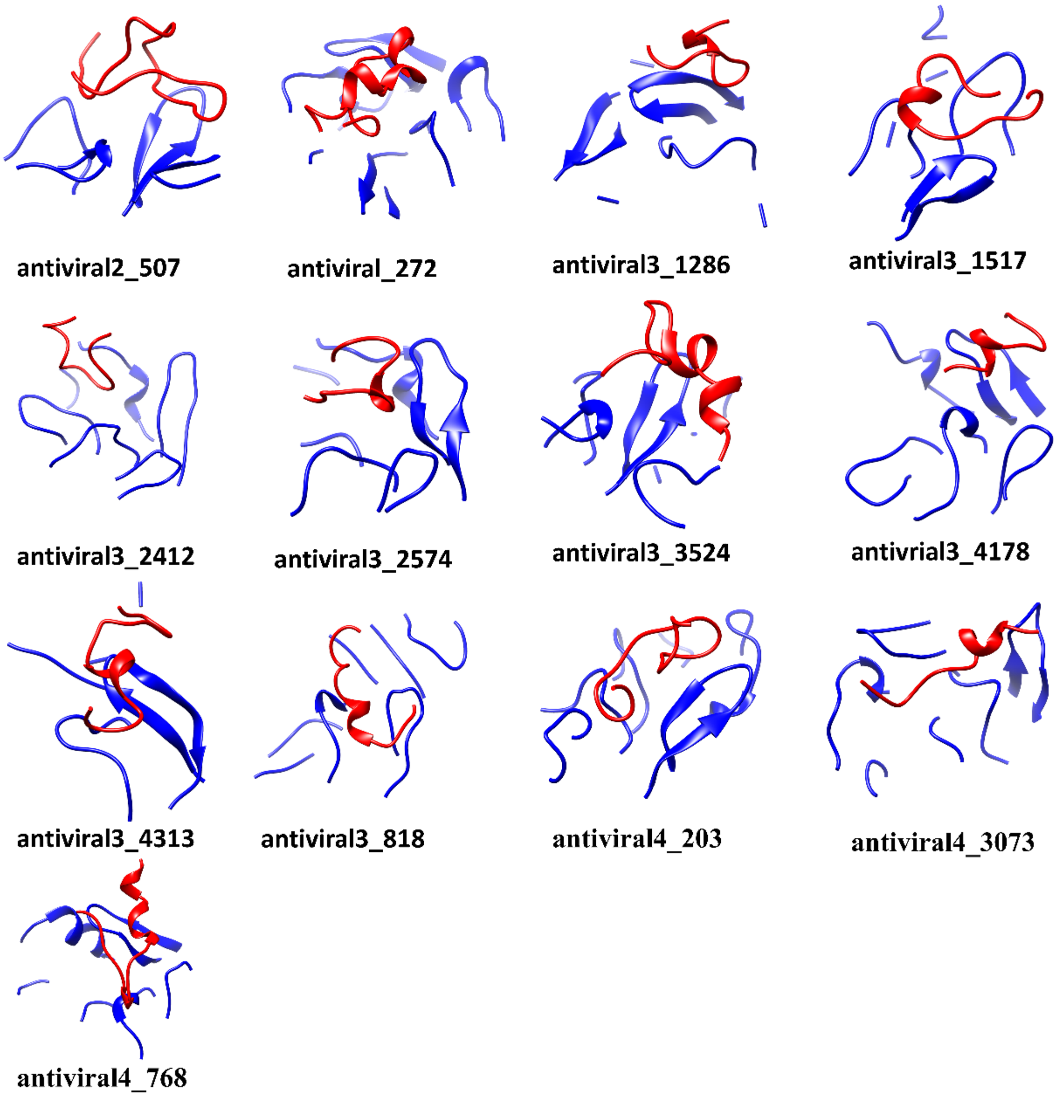


**Figure S12. The snapshot of the 13 final selected peptides with their binding protein pocket from the last frame of MD simulation.**

**Supplementary Table:**

**Table S1**. After pocket MD simulation, close contact residue pairs within 2.5Å between protein (Chain A) and peptide (Chain B).

| Name | Close contact residue pairs |
|---|---|
| antiviral_1443 | ASN119A PRO45B; THR21A ILE42B; ASN72A ASN23B; LEU67A ILE42B; GLN19A TYR46B; VAL73A ASN27B; LEU141A ILE10B; GLN19A ILE42B; ASN65A LYS41B; GLN74A THR31B; THR26A PRO45B; LEU141A PHE14B; GLN74A ASN27B; LEU67A LEU28B; ACE140A LEU6B; THR21A LYS41B; GLN19A LEU24B; GLN69A LEU24B; ACE22A LYS41B; TYR118A VAL13B |
| antiviral_1712 | MET165A ALA44B; ASN142A ASP40B; GLN192A ALA44B; LEU27A MET2B; THR25A SER3B; SER121A LEU14B; ASN142A TRP10B; ASN119A LEU14B; PRO168A GLN45B; THR24A ARG7B; GLU166A GLN45B; THR25A MET2B; GLN189A ALA44B; THR25A ARG7B; ASN142A SER1B; SER123A SER18B; LEU141A TRP10B; GLY143A TRP10B; GLY143A MET2B; THR25A THR6B; LEU141A LEU32B; ASN142A THR6B |
| antiviral_1996 | PRO168A MET43B; ASN142A ALA13B; PRO168A ALA40B; ASN119A PHE6B; SER121A PHE6B; PRO168A ALA44B; SER121A ALA2B; GLU166A LYS17B; MET47A GLY49B; LEU141A ALA13B; LEU141A LYS17B; MET47A VAL45B; GLN189A ALA44B; SER123A LYS7B; HIE41A LYS48B; TYR118A ILE9B; ASN142A VAL47B; GLN189A GLU41B; ALA191A LYS37B; TYR118A LYS10B; LEU141A ASN14B; HIE163A VAL47B; HIE164A LYS48B; SER121A LYS3B; ALA191A GLU41B; LEU141A LYS10B; LEU141A VAL47B; PRO122A LYS3B |

| | |
|---|---|
| antiviral_3445 | GLN189A CYS8B; THR24A CYS4B; THR25A LYS5B; MET47A CYS8B; GLU166A MET7B; THR25A TRP6B; THR45A TRP6B; THR24A TRP6B; GLN189A LYS10B; PRO168A LYS10B; THR24A LYS5B; PRO168A TRP11B; GLY23A GLY1B; MET47A VAL42B; LEU48A CYS37B; GLN189A CYS37B; ASN142A LYS5B; MET47A CYS37B; GLN189A ASP41B |
| antiviral_430 | HIE41A GLY1B; PRO168A THR50B; MET165A LEU2B; MET165A GLY1B; GLU166A LYS5B; THR45A VAL40B; HIE164A GLY1B; PRO168A SER49B; SER46A VAL40B; ASN142A TRP6B; GLN189A VAL4B; LEU141A TRP6B |
| antiviral_4465 | LEU141A CYS36B; ASN142A ILE17B; THR25A ARG16B; ACE164A LEU20B; THR25A THR19B; THR45A THR19B; GLU166A ARG40B; THR26A ILE17B; MET165A GLY22B; ASN142A GLY37B; GLU166A ALA38B; ACE140A ARG40B; LEU141A ARG40B; THR24A TYR15B; THR24A ARG16B; ASN142A CYS18B; THR26A ARG16B; ASN142A CYS36B; ASN142A SER21B |
| antiviral_490 | THR25A ASN27B; SER123A TRP11B; THR26A GLN26B; THR26A TYR23B; LEU27A TYR23B; TYR118A TRP11B; TYR118A ARG19B; THR24A ASN27B; LEU141A TRP11B |

| | |
|---|---|
| antiviral_822 | ACE140A CYS43B; MET47A GLU38B; LEU141A PRO22B; THR26A PRO24B; GLN192A TYR42B; ASN142A THR20B; ASN119A PRO24B; ASN142A ARG40B; ASN142A PRO22B; SER139A ARG46B; GLU166A TYR42B; MET47A ASN41B; THR25A GLN39B; ASN142A ILE33B; LEU167A GLN45B; NHE140A CYS11B; GLU166A GLN45B; GLY143A PRO22B; THR24A GLN39B; THR45A GLN39B; PRO168A ARG46B; MET47A GLN39B; ACE140A ARG46B; SER123A PHE9B; GLU166A ARG46B; GLY143A LEU23B; LEU141A PHE9B; SER46A GLN39B; THR24A TYR25B; ASN142A PHE21B; LEU141A LEU14B; PRO168A LEU49B |
